# Thermodynamic stabilization of the von Willebrand Factor A1 domain due to loss-of-function disease-related mutations

**DOI:** 10.1101/2022.03.10.483861

**Authors:** Angélica Sandoval-Pérez, Valeria Mejía-Restrepo, Camilo Aponte-Santamaría

## Abstract

The von Willebrand disease (vWD) is the most common hereditary bleeding disorder, caused by defects of the von Willebrand Factor (vWF), a large extracellular protein in charge of adhering platelets at sites of vascular lesion. vWF carries out this essential homeostatic task, via the specific protein-protein interaction between the vWF A1 domain and the platelet receptor, the glycoprotein Ib alpha (GPIB*α*). Upon the vWF activation triggered by the shear of the flowing blood. The two naturally occurring mutations G1324A and G1324S at the A1 domain, near the GPIB*α* binding site, result in a dramatic decrease of platelets adhesion, a bleeding disorder classified as type 2M vWD. However, it remained unclear how these two supposedly minor modifications lead to this drastic phenotypic response. We addressed this question using a combination of equilibrium-molecular dynamics (MD) and non-equilibrium MD-based free energy simulations. Our data confirm that both mutations maintain the highly stable Rossmann fold of the vWF A1 domain. These mutations locally diminished the flexibility of the binding site to GPIB*α* and induced a conformational change that affected the nearby secondary structure elements. Furthermore, we observed two significant changes in the vWF A1 domain upon mutation, the global redistribution of the internal mechanical stress and the increased thermodynamic stability of the A1 domain. These observations are consistent with previously-reported mutation-augmented melting temperatures. Overall, our results support the idea of thermodynamic conformational restriction of A1— before the binding to GPIB*α*—as a crucial factor determining the loss-of-function of the G1324A(S) vWD mutants.

## Introduction

The von Willebrand factor (vWF) is a multimeric plasma protein responsible for the binding and aggregation of platelets to sites of vascular injury during the first line of response against bleeding, known as primary hemostasis.^1, 2^ VWF arranges into monomers composed by several protein domains: D’D3, A1, A2, A3, D4, C1 to C6, and CK (figure 1A). These domains interact with various biomolecular partners. Importantly, the A1 domain interacts with the platelet glycoprotein IB*α* (GPIB*α*), collagen with A1 and A3, integrin with C4, and the coagulation factor VIII with D’D3, among other interactions.^3, 4^ Two vWF monomers dimerize at the CT/CK terminal domains via disulfite-bonds.^5, 6^ Moreover, these dimers form polymers which are assembled, compacted and stored in the trans-Golgi.^6^ Upon release to the blood stream, vWF becomes sensitive to hydrodynamic shear due to its large size reached upon polymerization. Accordingly, vWF undergoes shear-stress-driven reversible transitions from a globular to a stretched conformation, causing the exposure of hidden binding sites and therefore its activation. These transitions occur at physiological shear-stresses typically found in venules and arteries of the order of 10 dyn/cm^2^ to 50 dyn/cm^2^.^2, 7, 8^ The activated vWF recruits platelets at sites of injury an thereby promotes the formation of plugs to stop bleeding (Figure 1A).^6, 9^

**Figure 1:**
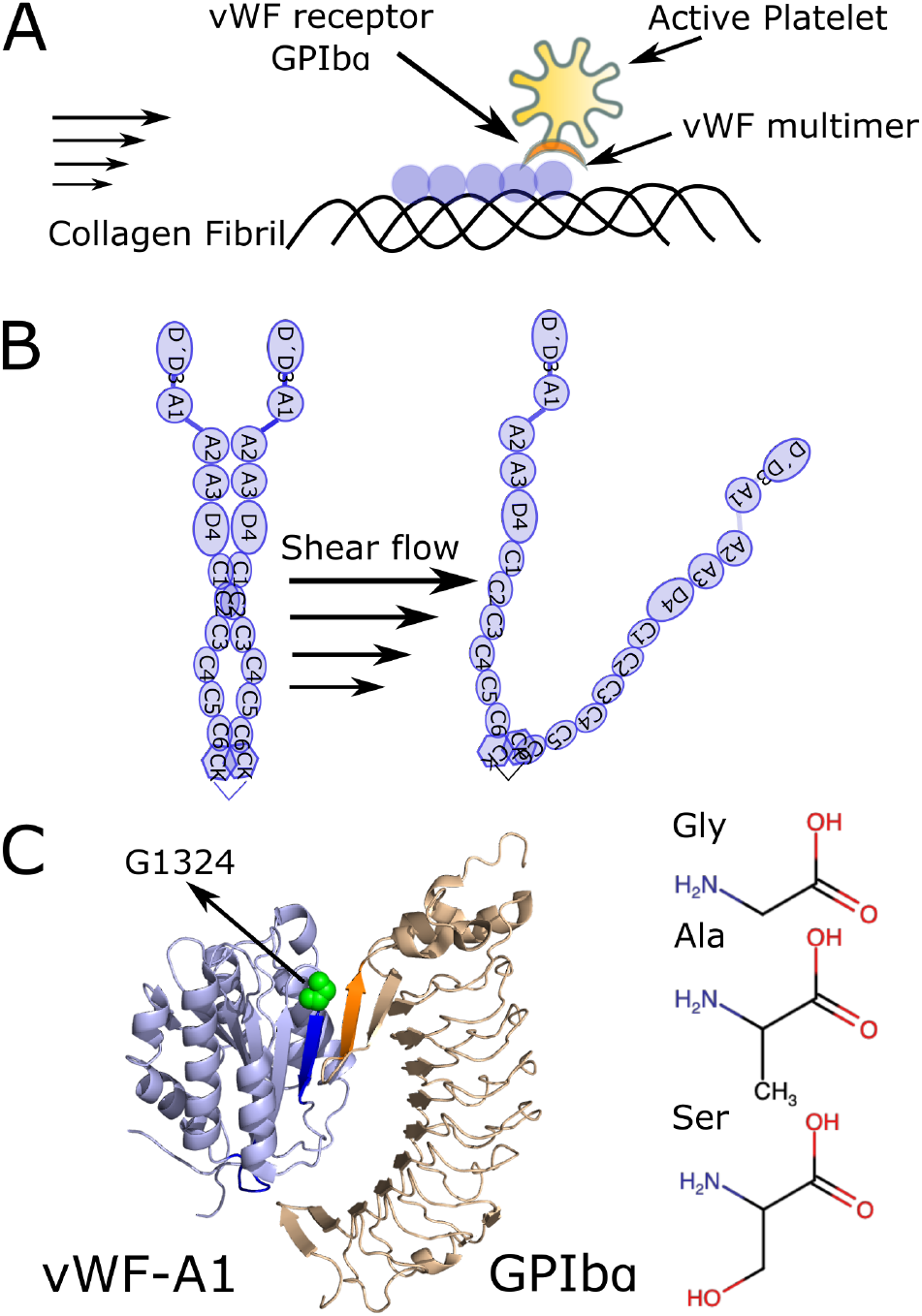
Scheme of von Willebrand factor (vWF) and type 2M disease mutations G1324A and G1324S. **A.** An injured blood vessel exposes its collagen fibrils. Triggered by the shear of the flowing blood, VWF unfolds to bind to the exposed fibrils and thereby recruit platelets at the site of injury. **B.** vWF is composed of several domains which are exposed by the action of the shear flow. Collagen binds to the A1 and A3 domains, whereas platelets bind to the A1 domain, via the interaction with the Glycoprotein IB*α* (GPIB*α*). **C.** *(Left)* VWF-A1 domain bound to the platelet receptor GPIB*α* is shown (binding sites highlighted in blue and orange, respectively). X-ray structure is shown (PDB id. 1SQ0^58^). The mutation site G1324 is highlighted in green. *(Right)* Chemical representation of the 1324 amino acid Gly and the vWD type 2M loss of function mutations Ser and Ala.

The malfunctioning of vWF causes von Willebrand disease (vWD), the most common inherited bleeding disorder with a prevalence of 0.6 to 1.3% in the population.^1, 10, 11^ The severity of the disease ranges from severe to mild bleeding.^1^ VWD is usually classified in three main types according to quantitative (Type 1 and 3) or qualitative (Type 2A, 2B, 2M and 2N) defects on vWF.^12^ Of particular interest are the qualitative defects increasing (2B) or decreasing (2M) the binding affinity to the platelet receptor GPIB*α*, without affecting blood levels of the vWF-multimers.

Two naturally occurring mutations, G1324A and G1324S (Figure 1C) are associated to the impairment of the vWF interaction platelet receptor GPIb*α*, causing vWD type 2M. These two mutations locate at the vWF A1, near the binding site of GPIB*α*. They significantly reduce vWF-platelet adhesion while maintaining the same vWF multimer level.^13^ Recent experiments have shown that the introduction of these mutations vastly increases the platelet-vWF dissociation rate and enhanced thermodynamic stability of the vWF A1 domain.^14^ Interestingly, the structure of the vWF-A1 G1324S mutant was not found to significantly deviate from the native A1 structure.^14^ Despite of the wealth of this information, the molecular mechanism by which these two mutations disrupt the vWF function are yet to be resolved.

The effect of naturally-inherited mutations on the activity of vWF has been intensively studied (for a comprehensive review see^15^). Quantitative characterization of the phenotypic response of VWD mutants is abundant, including defects in VWF expression and multimerization or the alterations in the binding of GPIB*α* and platelets. At the molecular level, X-ray crystallography revealed structural rearrangements in the A1 domain due to gain-of-function^14^ or loss-of-function^16^ mutations. Furthermore biochemical assays, e.g. by Tischer *et al.*,^14, 17, 18^ provided detailed thermodynamics and kinetic data regarding the interaction of A1 with GPIB*α* and its alteration due to mutations. Recently, force spectroscopy and platelet binding assays also demonstrated that two type gain-of-function vWD mutants alter vWF activity by destabilizing or disrupting the flanking auto-inhibitory modules of A1.^19–21^

Molecular dynamics (MD) simulations have also contributed enormously, not only to understand the functionality of several vWF domains, but also to the comprehension of vWF genetic disorders.^8, 21–35^ MD revealed key aspects of the interaction of A1 with GPIB*α*, ^28, 36^ the force-mediated auto-inhibition of this interaction imparted by the neighbor flanking regions,^22, 23, 27–29^ and the effect post-translational modifications has in controlling VWF-platelet interactions.^23, 37^ MD also characterized the interaction of DNA with the A1 domain, hindering the binding to platelets.^38^ In conjunction with free energy calculations, MD simulations also identified the mechanism of increased cleavage of the neighbor domain to A1, namely A2, due to an inherited disease-related mutation^24^ and the effect of methionine oxidation on the auto-inhibitory interaction between A1 and A2.^37^

Here, we use MD simulations and non-equilibrium MD-based free energy calculations to investigate the, so far lacking, molecular mechanism behind the loss-of-function vWD mutations G1324A and G1324S. Our simulations demonstrate that these two mutants alter the local flexibility of the mutation site near the GPIB*α* binding region, and more extensively influence the conformation of neighboring secondary elements near the binding site and globally redistribute the mechanical stress of the entire A1 domain. Free energy calculations show that these two mutants thermodynamically stabilize this domain, with the obtained free energy changes correlating very well with the urea-induced melting temperatures measured experimentally.^14^ Our data is thus consistent with a mechanism in which mutation-induced thermodynamic restriction of the A1 domain limits its binding to GPIB*α* and thereby reduces the binding of vWF to platelets.^14^ The used computational approach also constitutes the basis for the quantitative assessment of the thermodynamic stability of vWD-related A1 mutants.

## Materials and Methods

### Molecular Dynamics

MD simulations of the vWF A1 domain in isolation were conducted, starting from its experimentally determined structure (protein data bank code 1AUQ^39^). Three systems were considered: A1 in its wild type (WT) form and A1 with introduced mutations G1324A or G1324S. The mutations were introduced with Pymol.^40^ All the simulations were performed using the package GROMACS, version 2018.3,^41^ with the following combination of force fields: Amber99SB*ILDN for proteins,^42, 43^ TIP3P ^44^ model for water and Joung parameters for the ions.^45^ Sodium Chloride was added at a concentration of 0.15 M and counter ions were also added to neutralize the systems. The systems were minimized for 50000 steps. For each system, three replicas starting from the minimized conformation were generated and equilibrated for 10 ns of MD with position restraints on the heavy atoms of the protein (elastic constant of 1000kJ/*mol/nm*^2^). After equilibration, the position restrains were removed and 500 ns for each replica were performed for a total simulation time of 1.5*μ*s per system (4.5 *μ*s of cumulative unbiased simulation time). The temperature was kept at 310K, with velocity-rescaling thermostat,^46^ and time constant of 0.5 ps. Also, pressure was kept at 1 bar, with coupling constant of 1 ps, using the Berendsen barostat.^47^ The long range electrostatics were treated with the particle mesh Ewald (PME) technique.^48, 49^ Short-range interactions were considered through a Lennard-Jones potential within a cut-off distance of 1 nm. Bonds involving hydrogen atoms were constrained using the LINCS algorithm,^50^ and bonds and angles of water molecules were treated with SETTLE,^51^ hence allowing the integration of equations of motion at discrete time steps of 2 fs.

### Free Energy Calculations

MD-based free energy calculations were used to quantify the thermodynamic stability of the vWF A1 domain upon mutation. The free energy associated to the folding of the wild type A1 domain (Δ*G*_1_) and that corresponding to the mutated A1 domain, either G1324A or G1324S (Δ*G*_2_), were subtracted: ΔΔ*G* = Δ*G*_1_ – Δ*G*_2_. ΔΔ*G* was determined according to the thermodynamic cycle presented in Figure 5A, by computing the difference in free energy due to the mutation in the unfolded (Δ*G*_3_) and the folded (Δ*G*_4_) states: ΔΔ*G* = Δ*G*_1_ – Δ*G*_2_ = Δ*G*_3_ – Δ*G*_4_. Accordingly, a positive value of ΔΔ*G*, i.e. Δ*G*_1_ > Δ*G*_2_, implies that the mutant is thermodynamically more stable than the wild type A1 domain.

**Figure 2:**
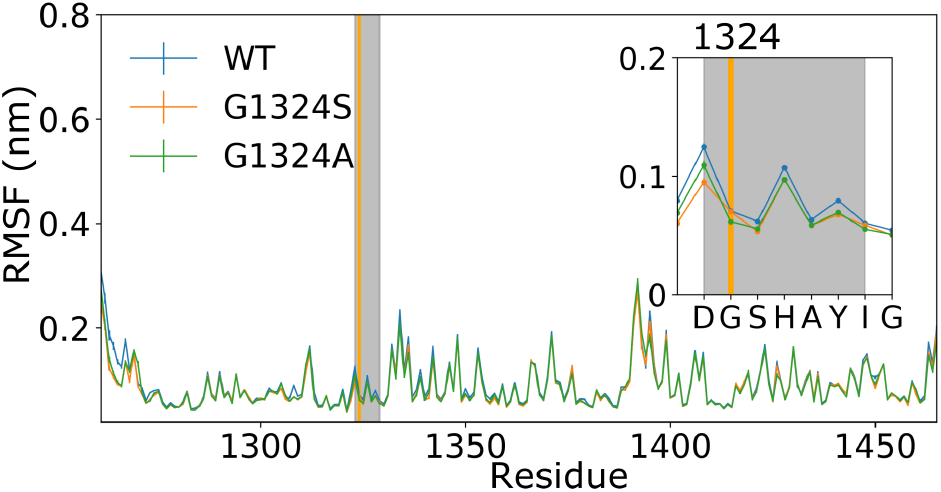
Root mean square fluctuations (RMSF) for each residue of the vWF A1 domain is presented for the wild-type WT (blue), G1324S (orange) and G1324A (green) mutants. The mutation point is specified with a vertical green line, and the binding site to GPIB*α* framed in a grey area. The *inset* is an enlarged view of the region including the binding site to GPIB*α* and 24 preceding residues.

**Figure 3:**
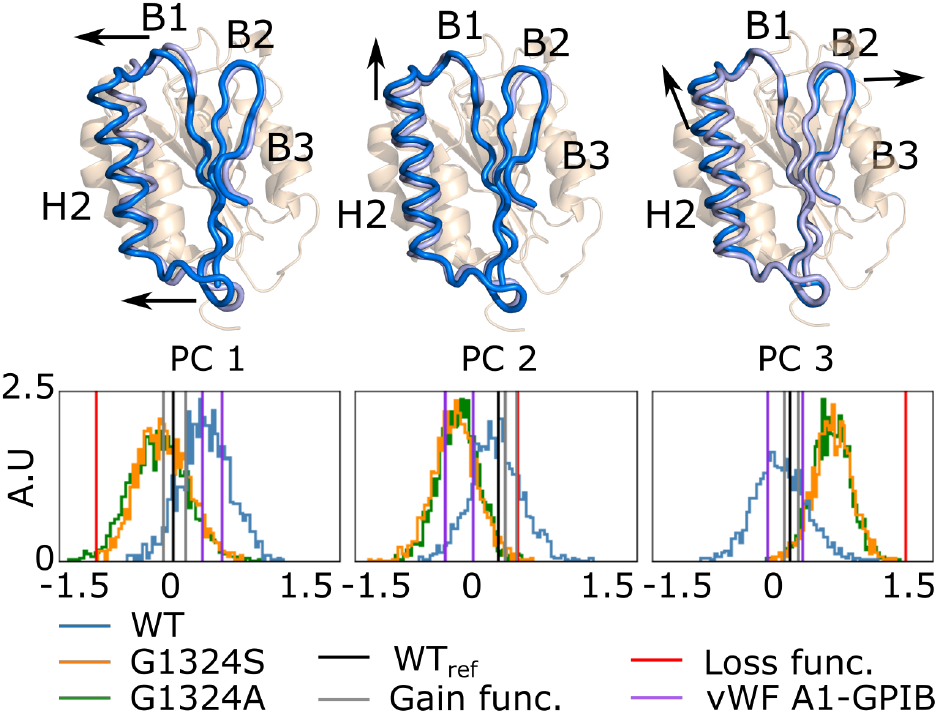
Principal component analysis of the vWF A1 domain. *(Up)* Three main collective movements of the vWF A1 domain retrieved by principal component analysis are considered. Residues 1284 to 1330 are highlighted in light and marine blue. The arrows indicate their movements with reference to the average structure depicted in wheat color in the background. *(Down)* Histogram of projections of MD trajectories onto the three principal components (PC 1, PC 2, and PC 3). WT, G1324A, and G1324S are represented in blue, green and orange respectively. Reported structures of the wild type vWF A1 domain (black); A1 mutants with either gain-of-function (grey) or loss-of-function (red), and A1 bound to GPIB*α*(purple) were also projected.

**Figure 4:**
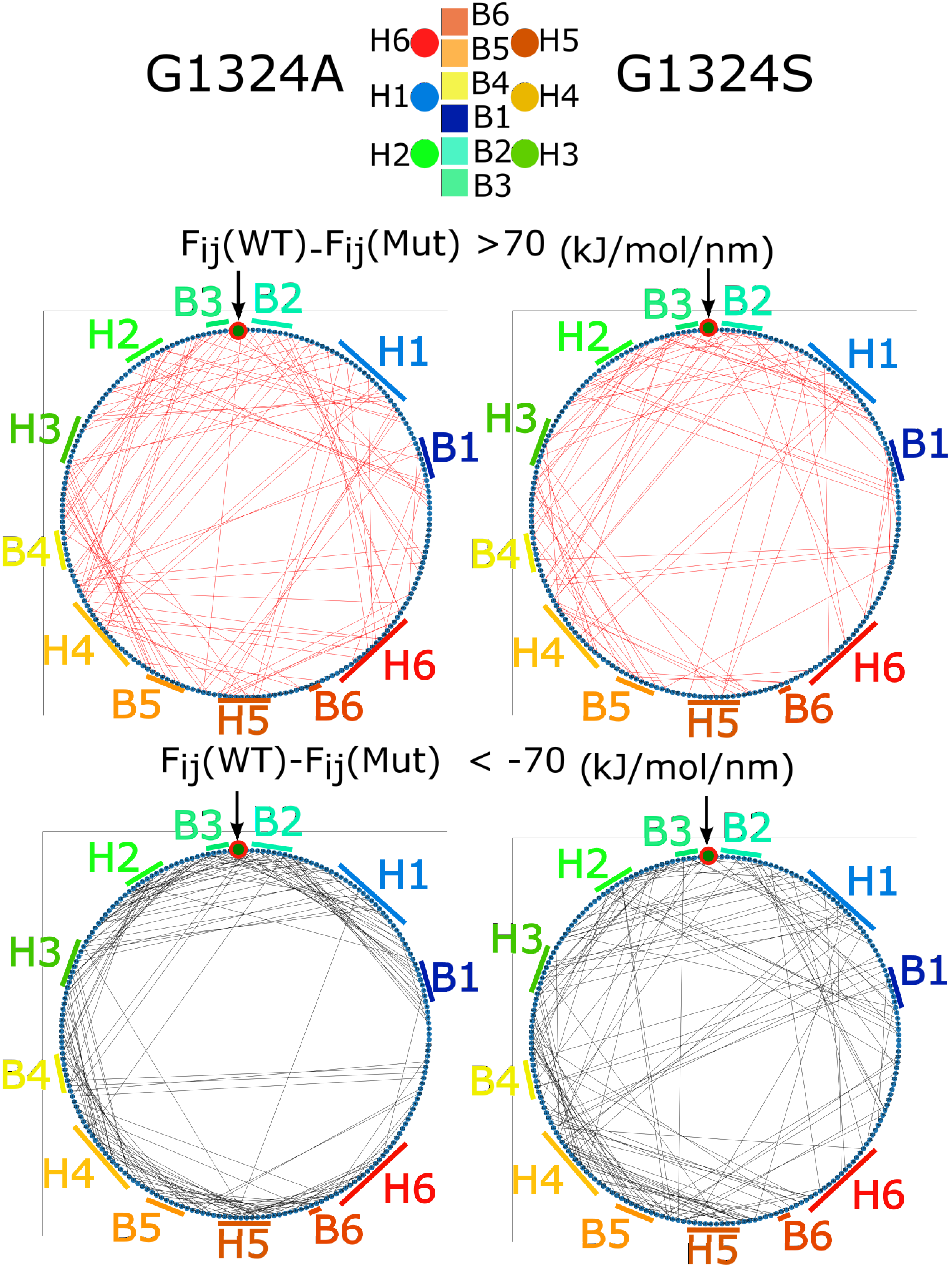
Force distribution analysis of the vWF A1 domain. The circle span the A1 sequence, from the N-terminus at 0 ° to the C-terminus at 360 °. Positions along the sequence of helices (H1 to H6) ad *β* strands (B1 to B6) are indicated. G1324 position is also highlighted with a red circle and and arrow. Pair-wise force for the mutants G1324A(left) and G1324S (right) minus that of the vWF A1 wild type (WT) domain is presented as lines connecting two points (two residues) of the circle (sequence). Pairs with a statistically significant difference (p-value< 0.01) and a force difference surpassing the threshold of 70 kJ/mol/nm were considered. Stronger pair-wise forces for the WT protein with respect to either mutant G1324A or G1324S (up) or vice versa (down) are presented.

**Figure 5:**
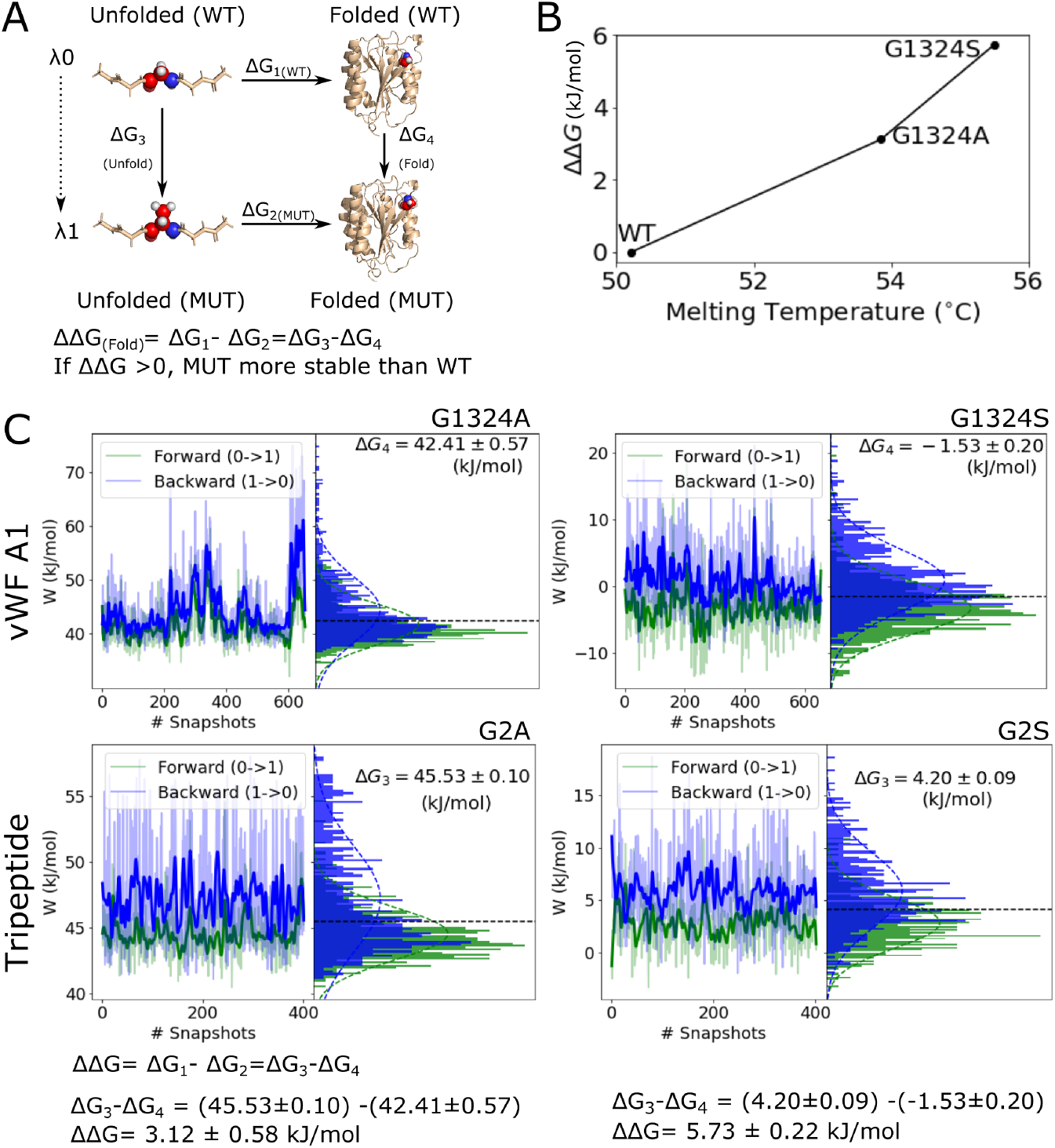
Thermodynamic stability of the vWF A1 domain analyzed by free energy calculations. **A.** Thermodynamic cycle used to assess the thermodynamic stability of the vWF A1 domain upon mutations. The horizontal arrows depict the folding process of the vWF A1 and the associated free energy Δ*G* in that process: wild type (up: WT; Δ*G*_1_) and the mutants G1324A or G1324S (down: MUT; Δ*G*_2_). The vertical arrows depict the alchemical transformation from the WT residue (glycine) to the mutated residue (either alanine or serine), for an unfolded tripeptide GXG (X=Gly, Ala or Ser), mimicking the unfolded state of the A1 domain (left; Δ*G*_3_), and for the folded vWF A1 (right; Δ*G*_4_). The indicated ΔΔ*G* is evaluated. Positive ΔΔG values indicate that the mutant is thermodynamically more stable than the wild type A1 domain. **B.** Calculated ΔΔ*G* as function of the experimental urea-induced melting temperature from^14^ is presented. **C.** Non-equilibrium Work recovered from thermodynamic transformation of the tripeptide GXG (bottom panels) or the vWF A1 domain (top panels) from the wild-type residue glycine to the mutated forms, either alanine (left) or serine (right). Work values are indicated for the forward transition, glycine to mutant (green) and minus work values for the backward transformation, mutant to glycine (blue). Work values are shown in the left panels and the resulting work distributions from them in the right panels. The intercept of the distributions (horizontal dashed line) corresponds to the free energy change for the folded protein Δ*G*_4_ or the tripeptide Δ*G*_3_. ΔΔ*G* is evaluated as the difference Δ*G*_3_ –Δ*G*_4_.

Thermodynamic integration was used to transition from the wild type (WT) to the mutant (MUT) states. A variable *λ* couples the Hamiltonian of the two states, such that *H*(*λ* = 0) = *H*_*WT*_ and *H*(*λ* = 1) = *H*_*MUT*_.^52^ The work associated to the transition is computed as 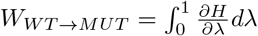. Subsequently, the work distribution associated to the forward transition WT→MUT (*P*_*WT→MUT*_(*W*)) and the reverse transition MUT→WT (*P*_*MUT→WT*_(*W*)) were computed to thereby calculate the free energy Δ*G* by using the Crooks fluctuation theorem:^53^

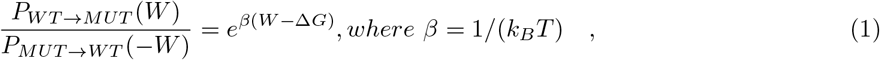

with *k*_*B*_ the Boltzmann constant, and *T* the temperature. Accordingly, the Bennett acceptance ratio (BAR) as a maximum likelihood estimator, proposed by *Shirts et al.*^54^ was used to derive the Δ*G* from the distribution of the non-equilibrium simulations. Assuming equal number of forward and reverse transitions, the BAR maximum-likelihood is expressed as follows:

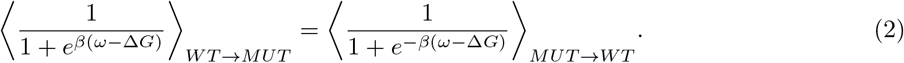

Here, ⟨⟩ denotes ensemble average. Finally, the uncertainty of Δ*G* was calculated by bootstrapping, solving equation (2) 100 times from randomly selected forward and reverse transitions.

600 conformations of folded vWF A1, obtained from the MD equilibrium simulations, served as starting structures for the non-equilibrium transitions. Hybrid topologies needed for thermodynamic integration were obtained with the *pmx* tool.^52^ The starting conformations were equilibrated for 1 ns at either *λ* = 0 or *λ* = 1 state. Transformations between *λ* = 0 to *λ* = 1 were of 400 ps in length. The end state of this simulations served as starting systems of an equilibration stept of 1 ns without position restraints and the further backwards transition. Similarly, three replicas of the tripeptide GXG (with X corresponding to Gly, Ala or Ser) were considered as representatives of the unfolded state. The use of tripeptides as reference to assess the thermodynamic stability of protein domains is common practice and has given very reasonable results in the past.^55^ 400 conformations were taken from 500-ns equilibrium MD simulations as starting positions for the forwards and backwards calculations. Non-equilibrium free energy simulations were performed with the GROMACS 4.6 version with an integrated soft-core potential function.^56^ The final free energies and their associated uncertainty were estimated using the *pmx* implementation.^57^

### Molecular Dynamics Analysis

Root mean square fluctuations (RMSF) were sampled every 10 ns (153 values per residue) to calculate average and standard mean errors. Principal component analysis (PCA) was used to quantify global changes in the conformation of the A1 domain. The covariance matrix was computed with the trajectories of the three replicas of the WT domain, considering the residues 1278 to 1333, which encompass the binding site to GPIB-*α* plus the neighboring secondary structure elements. The space defined by the PCA eigenvectors was used to project the corresponding trajectories of the wild-type protein and of the mutants G1324A and G1324S too. Additionally, experimentally reported structures of vWF A1 domains, containing loss-of-function (PDB id. 5BV8),^14^ or gain of function (PDB id. 4C29 and 4C2A) mutations,^16^ and the wild type vWF A1 domain bound to the GPIB*α* receptor (PDB id. 1SQ0)^58^ were also projected over the same PCA space. Force distribution analysis^59^ was used to compute inter-residue pair-wise forces (every 10 ns for a total of 150 uncorrelated data points). The difference in the average pair-wise force between the mutants (G1324A(S): MUT) and the wild type systems were computed for each residue pair (*i, j*) (Δ*F* (*i, j*) = *F*_*WT*_ (*i, j*) – *F*_*MUT*_ (*i, j*)). Accordingly, statistically significant pair-wise forces were presented (significance determined by a t-test, considering the N=150 uncorrelated data points).

## Results

### vWD type 2M mutations G1324A(S) diminished amino acid flexibility but only locally

We first monitored local changes on flexibility of the the vWF A1 domain due to the mutations G1324A and G1324S, in unbiased MD-simulations. To this end, we assessed the root mean square fluctuation (RMSF) for each residue of the protein (Figure 2). The RMSF of isolated vWF A1 domain confirmed the highly rigid structure of the A1 domain, imparted by its Rossmann fold. The core beta sheet surrounded by the helical elements displayed RMSFs below 0.1 nm. Moreover, the connecting loops also displayed reduced flexibility (RMSFs< 0.2 nm). Accordingly, the flexible termini presented large RMSF values. When focusing on the region near the mutation G1324, a reduction in RMSF of residues H1322 and D1323 was observed for the mutants, having G1324S the more marked effect (see inset of Figure 2). Except from this very localized change, near the mutation site, the mutations hardly changed the RMSF-per-residue pattern (Figure 2). Thus, mutations G1324A and G1324S modify the amino acid positional fluctuations but only locally near the mutation site.

### vWD type 2M mutations G1324A(S) changed conformation of a broad region covering the GPIB*α* binding site

We next checked if the mutants had a more global effect on the conformation of the vWF A1 domain. Accordingly, PCA was performed on the positions of residues 1284 to 1330, a region which covers the B1, B2, and B3 strands, along with the *α*-helix H2 (Figure 3, top). This region covers the immediate vicinity of the mutation site, importantly the GPIB*α* binding site at the strand B3. The first three principal components (PC) accounted for 33.84% of the total positional variance of the selected region (The collective movements associated with them are depicted in the upper section of Figure 3). The movements associated with PC 1 and 3 showed a translation of helix H2 apart from B1, thereby pulling with it the loop H1-B1. Interestingly, this loop that has been reported as key part for the binding to GPIB*α*.^58^ The MD trajectories were projected onto these three PCs and the distributions of these projections were plot separately for the WT and the mutant cases in Figure 3. The conformations explored by the mutants projected on this reduced PC space showed a high overlap between them, but only a partial overlap with the conformations sampled by the wild type counterpart (compare orange and green distributions with blue histogram in Figure 3). Consistently, projections of experimentally determined structures of vWF A1 WT (black),^39^ gain-of-function mutants (grey),^16^ and vWF A1 bound to the platelet receptor GPIB*α* (purple)^58^ showed a shared conformational space with the simulated vWF WT A1 domain. This is contrary to the loss-of-function mutant G1324S,^14^ which was shifted towards the conformational space explored by the mutants G1324A(S) in the PC 1 and 3 (Figure 3). Hence, the conformation of the region encompassing the GPIB*α* binding site was globally changed by the two vWD type 2M mutations.

### Mechanical stress redistribution caused by type 2M mutations G1324A(S)

Beyond structural changes highlighted in previous sections, we analyzed the redistribution in mechanical stress suffered by the vWF A1 domain due to the G1324A(S) mutations. For this, we used force distribution analysis (FDA).^59^ With FDA, we retrieved the difference in pair-wise force of the mutants minus that of the WT protein, for each residue pair (*i, j*), i.e. Δ*F* (*i, j*) = *F*_*WT*_ (*i, j*) – *F*_*MUT*_ (*i, j*). Accordingly, positive Δ*F* (*i, j*) indicates increased pair-wise forces between residue pairs in the wild type domain, while negative Δ*F* (*i, j*) indicates reduced pair-wise forces for the mutants. Figure 4 shows the pair-wise forces upon mutation, which hints to a global redistribution in mechanical stress compromising neighboring *β*-strands B1, B2, B3, and *α*-helices H2, but even reach more distant parts from the mutated point such as helices H4 and H6. The protein segments involved in force redistribution upon mutation thus extend to a broad region of the protein, beyond the point mutations site.

### G1324A and G1324S mutants increase thermodynamic stability of the vWF A1 domain

After examination of the changes in structure, dynamics and mechanical stress due to the G1324A and G1324S mutants on A1, we conducted free energy calculations to quantify the impact of these mutations on the thermodynamic stability of this domain. Such calculation was possible by considering the thermodynamic cycle presented in Figure 5A. This cycle allowed to compare the folding free energy of the vWF A1 mutant (Δ*G*2) with that for the wild type protein (Δ*G*1), by computing the energy required to mutate the wild-type vWF A1 domain, in either its unfolded ( Δ*G*_3_) or folded ( Δ*G*_4_) state. We used thermodynamic integration to extract the non-equilibrium work associated to the wild-type to the mutant transition (or vice versa) for the folded domain in 600 transitions (work distributions in Figure 5C) and for a tripeptide GGG representing the unfolded vWF A1 WT protein in 400 transitions (work distributions in Figure 5C). By using BAR suited for non-equilibrium free energy calculations,^54^ implemented in pmx,^57^ it was possible to estimate the free energy combining both forward and reverse work distributions. The absolute values obtained for Δ*G*_3_ compare well with the pre-computed values from the pmx data base, of 47.32 ± 0.25 kJ/mol for glycine to alanine and 7.46 ± 0.31 kJ/mol for glycine to serine, using same force-field as that of our calculations,^57^ thus validating our free-energy calculation protocol. The results obtained for the folded case, i.e. of Δ*G*_4_, are shown in the Figure 5C). The calculation of ΔΔ*G*, by subtracting Δ*G*_4_ from Δ*G*_3_, yielded 3.12 ± 0.58 kJ/mol for G1324A and 5.73 ± 0.22 kJ/mol for G1324S. Both are positive ΔΔ*G* values indicating that both mutants thermodynamically stabilize the vWF A1 domain. Our estimates correlate well with the urea-induced melting temperatures obtained previously experimentally ^14^ (Figure 5B).

## Discussion

We here studied the naturally-occurring mutants G1324A and G1324S associated with the vWD type 2M. By a combination of equilibrium MD simulations and non-equilibrium free energy calculations we assessed the changes in structure, dynamics, mechanical stress, and thermodynamic stability of the vWF A1 domain due to these mutations.

The vWF A1 domain was found very robust and maintained its structural stability despite of the mutations as expected from its Rossman Fold. The mutations G1324A and G1324S restricted the residue flexibility, but only locally for two residues near the mutation site, while the fluctuations of the rest of the protein remained largely unaltered (Figure 2). Nevertheless, PCA, of a region including the GPIB*α* binding site, i.e. strand B3 and loop H1-B2, and neighbor *β*-strands B1 and B2 and *α*-helix H2, showed a shift between the conformations explored by both mutants and the wild type protein (Figure 3). These conformational changes were accompanied by a mechanical-stress redistribution upon mutation of G1324 which extended way beyond the mutation point (Figure 4). Experimentally reported gain-of-function mutations of the vWF A1^16^ or variants bound to the platelet receptor GPIB*α* with high-affinity ^58^ coincided with the conformations explored by the vWF A1 wild type, while the low affinity variants^17^ towards conformations explored exclusively by the mutants (Figure 2). Thus, our observations support the idea that the here studied loss-of-function mutations alter the conformation and internal mechanical stress of a large portion of the vWF A1 domain and that this alteration occurs prior the binding to GPIB*α*.

The structural and mechanical changes observed here due to the introduction of the mutations could be also related to either over- or under-stabilization of the A1 domain. To test which of the two was the case, we assessed quantitatively the thermodynamic stability of the vWF A1 domain upon mutations, by using MD-based free energy calculations. The thermostability of the A1 domain was analyzed in terms of the unfolding process described in Figure 5. Our calculations yielded changes in the unfolding free energy ΔΔG of 3.12 ± 0.58 kJ/mol for G1324A and 5.73 ± 0.22 kJ/mol for G1324S (Figure 5C). Hence, our calculations demonstrate that both mutations G1324A and G1324S thermodynamically stabilize the A1 domain, with the latter one displaying the largest perturbation. Our results correlate very well with the experimentally reported urea-induced melting temperatures for the indicated mutants^14^ (Figure 5B). Overall, our study supports the mechanism proposed by Tischer et al.^14^ of mutation-induced thermodynamic conformational restriction of A1 limiting its binding to GPIB*α*, as the molecular mechanism governing the impaired function of these two vWD type 2M mutants.

The computational approach employed here appears very useful to systematically study the effect of vWD related mutations.^60^ In particular, the A1 domain appears as an excellent candidate for an *in silico* mutational scan, similar to that previously done for other proteins,^55, 61, 62^ but here aiming at assessing the connection between thermodynamics stability of the A1 domain and its main function of binding to the GPIB*α* platelet receptor.

## Conclusion

In summary, here we have computationally studied two naturally-occurring mutations of the von Willebrand factor that reduce its ability to bind platelets. Our calculations demonstrate that the two mutants thermodynamically stabilize the vWF A1 domain restraining it in a reduced sub-optimal conformational state for the binding to the GPIB*α* receptor. It will be highly interesting to exploit the computational approach used here in future studies to understand the effect of the other many vWD related mutations.

## Author Contributions

AS-P carried out the MD-Free energy calculations and analyzed the simulation data. VM-R carried out equilibrium MD simulations and analyzed the data. C-AS designed the study. AS-P and C-AS wrote the manuscript.

## Acknowledgments

The authors thank funding from the Max Planck tandem initiative of the University of Los Andes. Computing time was allocated at the Max Planck Computing and Data Facility in Garching, Germany, and the high-performance computing center of the University of Los Andes in Bogota, Colombia.

## References

1. Leebeek FW, Eikenboom JC. Von Willebrand’s disease. N. Engl. J. Med.. 2016;375(21):2067–2080.

2. Reininger AJ. Function of von Willebrand factor in haemostasis and thrombosis. Haemophilia. 2008;14:11–24.

3. Zhou YF, Eng ET, Zhu J, Lu C, Walz T, Springer TA. Sequence and structure relationships within von Willebrand factor. Blood. 2012;120(2):449–458.

4. Randi AM, Smith KE, Castaman G. von Willebrand factor regulation of blood vessel formation. Blood. 2018;132(2):132–140.

5. Katsumi A, Tuley EA, Bodo I, Sadler JE. Localization of disulfide bonds in the cystine knot domain of human von Willebrand factor. J. Biol. Chem. 2000;275(33):25585–25594.

6. Springer T. Von Willebrand factor, Jedi knight of the bloodstream. Blood. 2014:1412–1425.

7. Chiu J, Chien S. Effects of disturbed flow on vascular endothelium: pathophysiological basis and clinical perspectives. Physiol. Rev. 2011;91(1):327–387.

8. Grässle S, Huck V, Pappelbaum K, et al. von Willebrand factor directly interacts with DNA from neutrophil extracellular traps. Arter. Thromb. Vasc. Biol. 2014:1382–1389.

9. Rick M, Konkle B. Chapter 7: von Willebrand Disease. in Consultative Hemostasis and Thrombosis (Kitchens CS, Kessler CM, Konkle BA., eds.):90–102 Third Edition ed. 2013.

10. Schneppenheim R, Budde U. von Willebrand factor: the complex molecular genetics of a multidomain and multifunctional protein. J. Thromb. Haemost. 2011;9:209–215.

11. Huck V, Schneider MF, Gorzelanny C, Schneider SW. The various states of von Willebrand factor and their function in physiology and pathophysiology. Thromb. Haemost. 2014;111(04):598–609.

12. Sadler JE, Budde U, Eikenboom JCJ, et al. Update on the pathophysiology and classification of von Willebrand disease: a report of the Subcommittee on von Willebrand Factor.. J. Thromb. Haemost. 2006;4(10):2103–2114.

13. James PD, Goodeve AC. von Willebrand disease. Genet. Med. 2011;13(5):365–376.

14. Tischer A, Campbell JC, Machha VR, et al. Mutational constraints on local unfolding inhibit the rheological adaptation of von willebrand factor. J. Biol. Chem. 2016;291(8):3848–3859.

15. Budde U, Schneppenheim R, Eikenboom J, et al. Detailed von Willebrand factor multimer analysis in patients with von Willebrand disease in the European study, molecular and clinical markers for the diagnosis and management of type 1 von Willebrand disease (MCMDM-1VWD). J. Thromb. Haemost. 2008;6(5):762–771.

16. Blenner MA, Dong X, Springer TA. Structural basis of regulation of von Willebrand factor binding to glycoprotein Ib. J. Biol. Chem. 2014;289(9):5565–5579.

17. Tischer A, Machha VR, Frontroth JP, et al. Enhanced local disorder in a clinically elusive von Willebrand factor provokes high-affinity platelet clumping. J. Mol. Biol. 2017;429(14):2161–2177.

18. Tischer A, Brehm MA, Machha VR, et al. Evidence for the misfolding of the a1 domain within multimeric von Willebrand factor in type 2 von Willebrand disease. J. Mol. Biol. 2020;432(2):305–323.

19. Arce NA, Cao W, Brown AK, et al. Activation of von Willebrand factor via mechanical unfolding of its discontinuous autoinhibitory module. Nat. Commun. 2021;12(1):1–14.

20. Ju L, Dong J, Cruz MA, Zhu C. The N-terminal flanking region of the A1 domain regulates the force-dependent binding of von Willebrand factor to platelet glycoprotein Ib*α*. J. Biol. Chem. 2013;288(45):32289–32301.

21. Yago T, Lou J, Wu T, et al. Platelet glycoprotein Ib*α* forms catch bonds with human WT vWF but not with type 2B von Willebrand disease vWF. J. Clin. Investig. 2008;118(9):3195–3207.

22. Aponte-Santamaría C, Huck V, Posch S, et al. Force-sensitive autoinhibition of the von willebrand factor is mediated by interdomain interactions. Biophys. J. 2015;108(9):2312–2321.

23. Butera D, Passam F, Ju L, et al. Autoregulation of von Willebrand factor function by a disulfide bond switch. Sci. Adv.. 2018;4(2):eaaq1477.

24. Aponte-Santamaría C, Lippok S, Mittag JJ, et al. Mutation G1629E Increases von Willebrand Factor Cleavage via a Cooperative Destabilization Mechanism. Biophys. J. 2017;112(1):57–65.

25. Baldauf C, Schneppenheim R, Stacklies W, et al. Shear-induced unfolding activates von Willebrand factor A2 domain for proteolysis. J. Thromb. Haemost. 2009;7(12):2096–2105.

26. Brehm MA, Huck V, Aponte-Santamaría C, et al. von Willebrand disease type 2A phenotypes IIC, IID and IIE: A day in the life of shear-stressed mutant von Willebrand factor. Thromb. Haemost. 2014;112(7):96–108.

27. Posch S, Aponte-Santamaría C, Schwarzl R, et al. Mutual A domain interactions in the force sensing protein von Willebrand factor. J. Struct. Biol. 2017;197(1):57–64.

28. Interlandi G, Thomas W. The catch bond mechanism between von Willebrand factor and platelet surface receptors investigated by molecular dynamics simulations. Proteins. 2010;78(11):2506–2522.

29. Interlandi G, Ling M, Tu A, Chung DW, Thomas WE. Structural basis of type 2A von Willebrand disease investigated by molecular dynamics simulations and experiments. PloS One. 2012;7(10):e45207.

30. Interlandi G, Yakovenko O, Tu A, et al. Specific electrostatic interactions between charged amino acid residues regulate binding of von Willebrand factor to blood platelets. J. Biol. Chem. 2017;292(45):18608–18617.

31. Goto S, Oka H, Ayabe K, et al. Prediction of binding characteristics between von Willebrand factor and platelet glycoprotein Ib*α* with various mutations by molecular dynamic simulation. Thromb. Res. 2019;184:129–135.

32. Chen Z, Lou J, Zhu C, Schulten K. Flow-induced structural transition in the *β*-switch region of glycoprotein Ib. Biophys. J. 2008;95(3):1303–1313.

33. Rana G, Pathak RK, Shukla R, Baunthiyal M. In silico identification of mimicking molecule (s) triggering von Willebrand factor in human: a molecular drug target for regulating coagulation pathway. J. Biomol. Struct. Dyn. 2020;38(1):124–136.

34. Kuzmanic A, Pritchard RB, Hansen DF, Gervasio FL. Importance of the force field choice in capturing functionally relevant dynamics in the von Willebrand factor. J. Phys Chem. Lett. 2019;10(8):1928–1934.

35. Shiozaki S, Takagi S, Goto S. Prediction of molecular interaction between platelet glycoprotein Ib*α* and von Willebrand factor using molecular dynamics simulations. J. Atheroscler. Thromb. 2015:32458.

36. Zou X, Liu Y, Chen Z, Cárdenas-Jirón GI, Schulten K. Flow-induced *β*-hairpin folding of the glycoprotein Ib*α β*-switch. Biophys. J. 2010;99(4):1182–1191.

37. Tsai R, Interlandi G. Oxidation shuts down an auto-inhibitory mechanism of von Willebrand factor. Proteins. 2021;89(6):731–741.

38. Sandoval-Pérez A, Berger RM, Garaizar A, et al. DNA binds to a specific site of the adhesive blood-protein von Willebrand factor guided by electrostatic interactions. Nucleic Acids Res. 2020;48(13):7333–7344.

39. Emsley J, Cruz M, Handin R, Liddington R. Crystal structure of the von Willebrand factor A1 domain and implications for the binding of platelet glycoprotein Ib. J. Biol. Chem. 1998;273:10396–10401.

40. DeLano WL. The PyMOL Molecular Graphics System, Version 2.3. 2020.

41. Abraham M, Spoel D, Lindahl E, Hess B, team.. Gromacs User Manual Version 2018.3. 2018.

42. Lindorff-Larsen K, Piana S, Palmo K, et al. Improved side-chain torsion potentials for the Amber ff99SB protein force field. Proteins. 2010;78(8):1950–1958.

43. Best RB, Hummer G. Optimized molecular dynamics force fields applied to the helix-coil transition of polypeptides. J. Phys Chem. B. 2009;113(26):9004–9015.

44. Jorgensen WL, Chandrasekhar J, Madura JD. Comparison of simple potential functions for simulating liquid water. J. Chem. Phys. 1983;79(2):926–935.

45. Joung I, Cheatham TE. Determination of alkali and halide monovalent ion parameters for use in explicitly solvated biomolecular simulations. J. Phys Chem. B. 2008;112(30):9020–9041.

46. Bussi G, Donadio D, Parrinello M. Canonical sampling through velocity rescaling. J. Chem. Phys. 2007;126(1):014101.

47. Berendsen HJ, Postma JP, Van Gunsteren WF, Dinola A, Haak JR. Molecular dynamics with coupling to an external bath. J. Chem. Phys. 1984;81(8):3684–3690.

48. Darden T, York D, Pedersen L. Particle mesh Ewald: An N log (N) method for Ewald sums in large systems. J. Chem. Phys. 1993;98(12):10089–10092.

49. Essmann U, Perera L, Berkowitz ML, Darden T, Lee H, Pedersen LG. A smooth particle mesh Ewald method. J. Chem. Phys. 1995;103(19):8577–8593.

50. Hess B, Bekker H, Berendsen HJC, Fraaije JGEM. LINCS: a linear constraint solver for molecular simulations. J. Comp. Chem. 1997;18(12):1463–1472.

51. Miyamoto S, Kollman PA. Settle: An analytical version of the SHAKE and RATTLE algorithm for rigid water models. J. Comp. Chem. 1992;13(8):952–962.

52. Gapsys V, Michielssens S, Peters JH, Groot BL, Leonov H. Calculation of binding free energies. Methods Mol. Biol. 2015;1215:173–209.

53. Crooks GE. Nonequilibrium Measurements of Free Energy Differences for Microscopically Reversible Markovian Systems. J. Stat. Phys. 1998;90(5):1481–1487.

54. Shirts MR, Bair E, Hooker G, Pande VS. Equilibrium free energies from nonequilibrium measurements using maximum-likelihood methods.. Phys. Rev. Lett. 2003;91(14):140601.

55. Seeliger D, De Groot BL. Protein thermostability calculations using alchemical free energy simulations. Biophys. J. 2010;98(10):2309–2316.

56. Gapsys V, Seeliger D, De Groot BL. New soft-core potential function for molecular dynamics based alchemical free energy calculations. J. Chem. Theory Comput. 2012;8(7):2373–2382.

57. Gapsys V, Michielssens S, Seeliger D, De Groot BL. pmx: Automated protein structure and topology generation for alchemical perturbations. J. Comput. Chem. 2015;36(5):348–354.

58. Dumas JJ, Kumar R, McDonagh T, et al. Crystal structure of the wild-type von Willebrand factor A1-glycoprotein Ib*α* complex reveals conformation differences with a complex bearing von Willebrand disease mutations. J. Biol. Chem. 2004;279(22):23327–34.

59. Costescu BI, Gräter F. Time-resolved force distribution analysis. BMC Biophys. 2013;5(1):5.

60. Jong A, Eikenboom J. Von Willebrand disease mutation spectrum and associated mutation mechanisms. Thromb. Res. 2017;159:65–75.

61. Gapsys V, Michielssens S, Seeliger D, Groot BL. Accurate and rigorous prediction of the changes in protein free energies in a large-scale mutation scan. Angew. Chem. Int. Ed. 2016;55(26):7364–7368.

62. Gapsys V, Pérez-Benito L, Aldeghi M, et al. Large scale relative protein ligand binding affinities using non-equilibrium alchemy. Chem. Sci. 2020;11(4):1140–1152.

